# Random walks with spatial and temporal resets may underlie searching movements in ants

**DOI:** 10.1101/2024.02.20.581181

**Authors:** Valentin Lecheval, Elva J.H. Robinson, Richard P. Mann

## Abstract

Many ant species are central place foragers, living in a nest and exploiting the surrounding environment. It is however unclear how their exploration behaviour relates to the emerging exploited area. Ants provide a great opportunity to study the emergence of foraging range from individual movements, given the potentially large number of scouting workers involved. Here, we introduce a random walk model with stochastic resetting to depict the movements of searching ants. Stochastic resetting refers to spatially reset at random times the position of agents to a given location, here the nest of searching ants. We investigate the effect of a range of resetting mechanisms by changing how the probability of returning to the nest depends on the duration of unsuccessful foraging trips. We compare the macroscopic predictions of our model to laboratory and field data. We find that the probability for searching ants to return to their nest decreases as the number of foraging trips increases, resulting in scouts going further away from the nest as the number of foraging trips increases. Our findings highlight the importance of resetting random walk models to depict the movements of central place foragers and nurtures novel questions regarding the searching behaviour of ants.

## 1 Introduction

Animals that carry resources back to a particular site such as their nest, are called central place foragers. The decisions of central place foragers are affected by external factors such as food quality or distance to food patches [1–4]. The mechanistic understanding of the movements of animals in general and of central place foragers in particular concerns the relation between the internal state of the animal, its locomotion and navigation abilities to the environmental factors (e.g. distribution of resources) [5, 6]. The feedback loop between food patch characteristics and the foraging behaviour, driven by the ability to locate food, results in animals exhibiting various movement patterns, including random walks [7, 8] and Lévy walks [9–11]. From the movement behaviour of animals (“microscopic scale”) emerges a foraging area or a territory (“macroscopic scale”), that can be mechanistically inferred from the movements and behaviours of individuals [12–14]. Such a mechanistic understanding of territory or home range formation in animals is instrumental to predict the impact of environmental perturbations on the survival and reproduction abilities of a species [14–16].

Social insects, including ants, are a great opportunity to study phenomena that emerge from self-organisation [17]. For instance, the potentially large number of workers in a social insect colony ensures that the macroscopic scale emerging from the multiple interactions at the microscopic scale has a spatial and temporal relevance to explain colony-level decisions and behaviours. It is particularly true for the emergence of a foraging area or a territory from individual movements [18, 19], such as in colonies of social bees, where the scouting movements of workers can be used to understand the complex dynamics between pollinators and plant reproduction [20]. Many ant species are central place foragers, living in a nest and exploiting the surrounding environment. It is still unclear however how the mechanisms of searching ants, including their movements, relates to the exploited foraging area of a colony. More specifically, a modelling framework unifying known behavioural mechanisms involved in searching for ants with a macroscopic description of the foraging area at the colony level is lacking.

The movements of single ants navigating a simple laboratory environment can be described in fine detail by a correlated random walk [21], although other movement types such as meandering may also be present [22]. At the individual scale, this behaviour results in a series of straight moves of varying lengths, separated by turning angles favouring new directions close to the previous ones, meaning that the ants have a tendency to make small deviations at each reorientation event. At the collective scale, the description of a population of ants moving with a correlated random walk corresponds to a diffusive pattern. For a given ant nest, if ants whose task is to search the surroundings to find resources – so-called scouts, move with a correlated random walk, this would result in all scouts, on average, progressively going away from their nest and their density decreasing over time in areas close to the nest. In other words, this would result in an unrealistic outcome of scouts moving away from the nest, presumably until they find resources, and the density of scouts being homogeneously very low around the nest. This is unrealistic because the time to find resources may exceed the time a scout can survive outside its nest, and the energy costs associated with such an exploration pattern would be very high at the population level. By investigating the behaviour of ants on their inbound journey, returning to their nest and searching for a nest entrance, studies have revealed a looping behaviour, where individuals go back and forth from their initial location, gradually increasing the sizes of the loops [23–26], in agreement with a looping behaviour reported in fire ants searching for food [27]. This searching behaviour of foraging ants may therefore well be involved in outbound journeys, when foragers are exploring the environment without preconceived ideas about potential favourable locations or when they are not involved in exploiting a food patch after recruitment [28]. This searching strategy involves sinuous random walk outbound movements and straighter trajectories back to the nest [28]. The searching behaviour of ants has been investigated theoretically with models developed at the individual scale, but with little consideration for the associated emerging collective pattern, namely the foraging area [e.g. 22, 23, 29, 30].

There are methods to estimate, in the field, the foraging area and territory of an ant colony, based on foraging or aggressiveness responses and nest dispersion [31–33]. Theoretical models, based on the same principles, have also been proposed to estimate the size and arrangements of the territories of ant populations [34, 35]. These methods, theoretical and empirical alike, often reveal, infer or predict the general shape of the borders of a territory (e.g. circular [31]) at a given time, but do not provide a mechanistic and dynamic description of its emergence [14]. It is, however, worth noting a few quantitative properties of searching and foraging ants at the macroscopic scale that help to characterise the foraging area. In ants searching for their nest entrance in an unknown environment, several studies on various species (the Australian desert ant *Melophorus bagoti* and a western Mediterranean ant *Aphaenogaster senilis*) have reported that the distribution of the distance between individuals and the origin where they started their search could be depicted by two exponential distributions [25, 28]. This suggests there is a general mechanism in searching ants moving around a central place that give rise to ants being exponentially distributed – therefore indicating that the probability to find resources decreases with the distance from the nest as an exponential. In fire ants, it was found that the time spent by scouts outside their nest before returning was distributed with a power law [27]. In other words, the probability for a single scout to return to the nest was not found to be constant in time: the longer the individual spends outside the nest, the lower the probability to return (compatible with the qualitative reports of increasing loops exhibited by searching desert ants).

In a recent study, we made the assumption that scouts have a tendency to return to their nest even if they are unsuccessful in finding resources and predicted that the resulting distribution of scouts around their nest would be exponential [4]. It has not, however, been thoroughly shown how the individual mechanisms involved in searching behaviours previously reported in the literature are connected with the macroscopic description and the range of the foraging area [36]. Here, we aim to develop a modelling framework that will unify both the individual and collective descriptions of scouting ants. We formulate a spatially-explicit model, extending an ant correlated random walk model with a tendency for ants to stochastically return to their nest. Such models, depicting random walk movements with stochastic returns to a central place have recently received attention in statistical physics, leading to an emerging literature on random walks with stochastic resetting [37]. This framework is well-suited to describe central place foraging [20, 38]: it allows description of the system at both microscopic and macroscopic scales and the analysis of the effects of varying both movement patterns and the dynamics of the returning (resetting) episodes [39, 40]. This class of models have already been used to investigate the foraging movements of honey bees [20, 38], where it was suggested that the rate of returning to the nest was constant in time [20], similar to what has been proposed in ants [4].

Here, we extend a random walk model for ant movements [21] with a stochastic reset process to evaluate whether or not the hypothesis of a nest-returning rate constant in time for unsuccessful foragers is compatible with empirical data at both the individual and collective levels. We specifically explore how constant, or decreasing linearly or exponentially rates to return to the nest with respect to time determine the duration of journeys outside the nest and the distribution of scout distances from their nest. We evaluate these alternatives by reanalysing an existing dataset of searching ants and using published results reported in various ant species, both in laboratory and field settings. We suggest important metrics to discriminate different models of random walks with resetting mechanisms and discuss the implications of our findings to understand how the exploration range of an ant colony is derived from simple movement rules and what may affect resets in scouting ants.

## 2 Material and Methods

### 2.1 Empirical data

In order to investigate the searching behaviour of scouting ants, we analyse data published in [41] with dataset available on Dryad ^1^. Data consist of 21 trajectories of single *Temnothorax albipennis* worker ants from 3 colonies (i.e. 7 individual per colony) exploring a square arena recorded by video and tracked for 45 min at 10 fps. In our analysis we consider all distances within 8.4 cm from the nest entrance as the nest, since this area is made of a distinct curvature and material compared to the rest of the arena [41]. We restrict our analysis to the data from the clean treatment, in which chemical markers of previously tested ants are removed between each trial.

For each ant trajectory, the moments when the individuals reach the nest are identified, leading to series of trajectories (scouting bouts) in which the ants are walking in the arena outside of their nest. The instants when the ants are the furthest away from their nest during these scouting bouts are characterised as U-turns, that is when the ants start to walk towards their nest (inbound journey). The time of the U-turns, distance from the nest at U-turns and the number of U-turns for each ant are recorded.

### 2.2 Model formulation

We use as a basis the correlated random model of the Boltzmann Walker which has already been used to describe the searching movements of single individuals in *Lasius niger* moving on tilted planes [21]. In this model, agents move straightforward at a constant speed, before turning to other directions, biased towards the preceding headings, effectively producing a trajectory made of a series of segments, separated by discrete reorientation events. The length of the segments is exponentially distributed, with an average length *λ* called the *mean free path*. The probability density function underlying the choices of new directions is symmetric around the incoming direction, and can be more or less concentrated around it. This concentration is quantified by a parameter *g*, the mean cosine of the orientation deviation, which indicates the heading persistence (from *g* = 0 for a complete reorientation process, or null persistence, to *g* = 1 for null deviations, or complete persistence). The two parameters *λ* and *g* are the two parameters of the Boltzmann Walker model.

We extend this model by adding a new parameter *p*(*r*), the probability to return to the nest at each reorientation event, with *r* the number of reorientation events experienced by the agent since its first departure from the nest. At each reorientation event, the position of the agent is reset to the nest with probability *p*(*r*). When the spatial reset occurs, the agents returns to its nest in a straight path (ballistic return), walking at a constant speed, and the next reorientation event takes place from the nest. The fact that inbound bouts are straighter than outbound ones was previously reported in *Aphaenogaster senilis* ants [28] and studied theoretically in the context of random walk with stochastic resetting [42]. We will explore three scenarios of how the rate of returns to the nest depends on the number of reorientation events in this study, with *p*(*r*) = *c*, constant with the number of reorientation events or with *p*(*r*) decreasing linearly or exponentially with the number of reorientation events (Figure 1).

**Figure 1.**
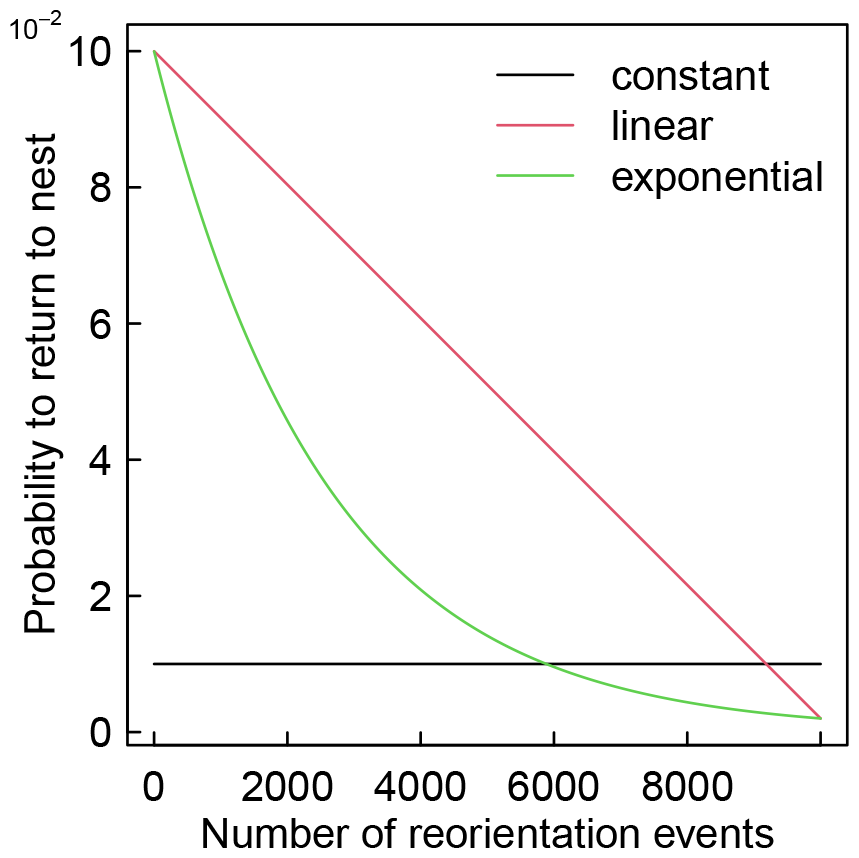
Three theoretical scenarios of how the probability to return to the nest depends on the number of reorientation events experienced by the agent.

The location of an ant at reorientation event *r* is updated as

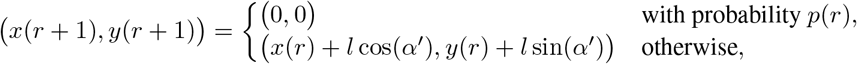

with *l* the length of the next path, sampled from an exponential distribution of rate 1*/λ* and *α*^*′*^ = *α* + *d*, the new heading of the agent, *d* being an angular deviation obtained from elliptical angular sampling (Algorithm 1).

#### Algorithm 1

Elliptical angular sampling. This function return an angular deviation given the average cosine of the deviation *g* [21]. 𝒰 (*a, b*) stands for the continuous uniform distribution with parameters *a* and *b*.

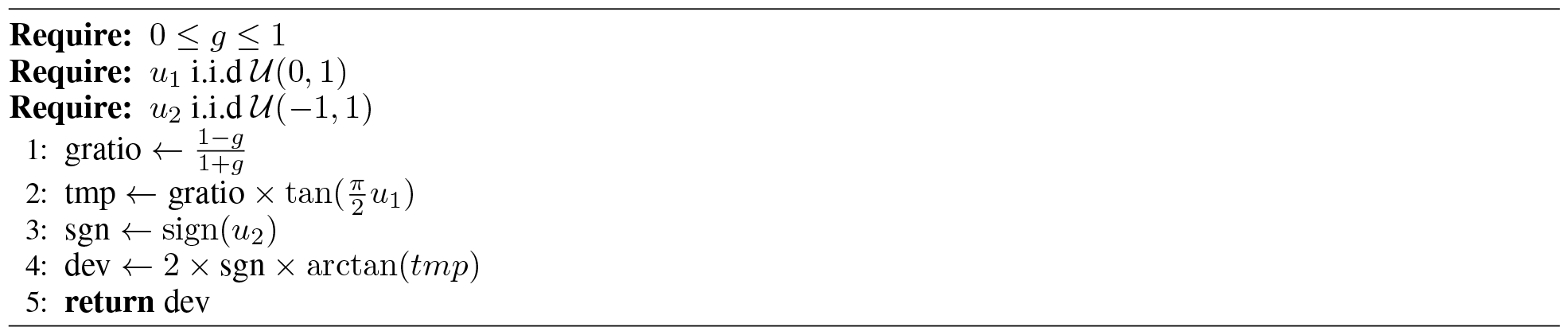

### 2.3 Model simulations

In all the simulations, agents start from the same initial location (*x, y*) = (0, 0), depicting the nest of the colony, and move with the same parameters of reference across all simulations, with the heading persistence *g* = 0.6, the mean free path *λ* = 0.01 m and the speed *s* = 0.04 m.*s*^*−*1^ [21].

#### 2.3.1 Distributions of distance to the nest and mean square displacement

The model is simulated with *n* = 20000 non-interacting searching agents moving for a given number of reorientation event *r* (effectively the simulation duration). In each treatment, all agents move for *r* reorientation events, starting from their nest at the beginning of each treatment. We set 6 treatments *r ∈* {50, 100, 500, 1000, 5000, 10000} reorientation events to estimate the distribution distances from the nest and 10 treatments *r ∈* {1, 10, 50, 100, 500, 1000, 2500, 5000, 7500, 10000} reorientation events to calculate the mean square displacement. The distance of the *n* = 20000 agents from the nest after *r* reorientation events is calculated for agent *i*,

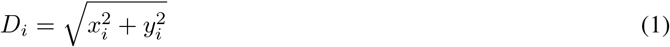

with (*x*_*i*_, *y*_*i*_) the coordinates of agent *i* after *r* reorientation events. The mean square displacement is calculated for each number of reorientation event *r* as

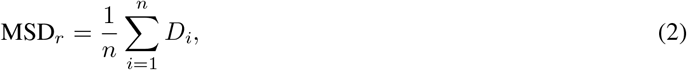

with (*x*_*i*_, *y*_*i*_) the coordinates of agent *i* after *r* reorientation events.

#### 2.3.2 Survival curves and maximum distance reached in outbound journeys

We ran simulations monitoring metrics during the entirety of the exploration journeys. These metrics, evaluated at each reorientation event of each agent, include (i) the time spent in an outbound journey, used to make survival analysis of the time spent outside of the nest exploring the environment and (ii) the maximum distance from the nest reached by agents during these outbound journeys. To keep track of the elapsed time, we used a constant speed for the agents, set to *s* = 0.04 m.s^*−*1^.

#### 2.3.3 Probability to find resources

We also ran simulations to calculate how the probability to find resources varies with distance to the nest for various rules of returning to the nest. We set 41 resources at a range of distances from the nest (every centimetre from 10 to 50 centimetres from the nest), with random directions sampled from a distribution uniform around the nest. 1000 agents, each with a constant number of reorientation events set to 500, have the opportunity to find each of the 41 resources in as many simulation runs. A resource is considered found if an agent passes within 5 cm of the resource. The probability of a resource being found is set as the proportion of agents which successfully found it within 500 reorientation events.

## 3 Results

### 3.1 Empirical data

We find that (i) ants are navigating back and forth to their nest (0.8±0.28 return events per ant per min, mean ±sd) and (ii) that this results in distance from the nest being exponentially distributed (Figure 2a-b). Figure 2b actually suggests the presence of two distinct exponential distributions, one at very short distances from the nest (less than 7 cm) and another one for greater distances. This seems to be the result of the distance distribution not being stationary in time: at the beginning of the observation period, ants stay within 20 cm of their nest, then they are progressively found further away as time goes – being exponentially distributed at each of these time periods. The distribution averaged over the entire observation window therefore reflects this tendency. To investigate further the individual mechanisms responsible for the exponential distribution of the distance of scouts to their nest, we record the time at which scouts are the furthest away from their nest for each scouting event. These times are then taken to be the times of decisions of scouts to return to their nests. We construct the survival curve of ants still on their outbound journey from these times (Figure 2c). We find that for short outbound journeys (less than 10 s), the survival curve is close to an exponential: the probability for ants to return to their nest in the next time interval is constant. For longer outbound journeys (more than 10 s), the survival curve follows a power law: the longer a worker remains scouting, the lower becomes the probability that it will return home in the next time interval. The scouting ants therefore go on average further away as the time since the beginning of the experiment started increases (Figure 2d), confirming what was found by looking at the distribution of distances at different time windows (Figure 2b).

**Figure 2.**
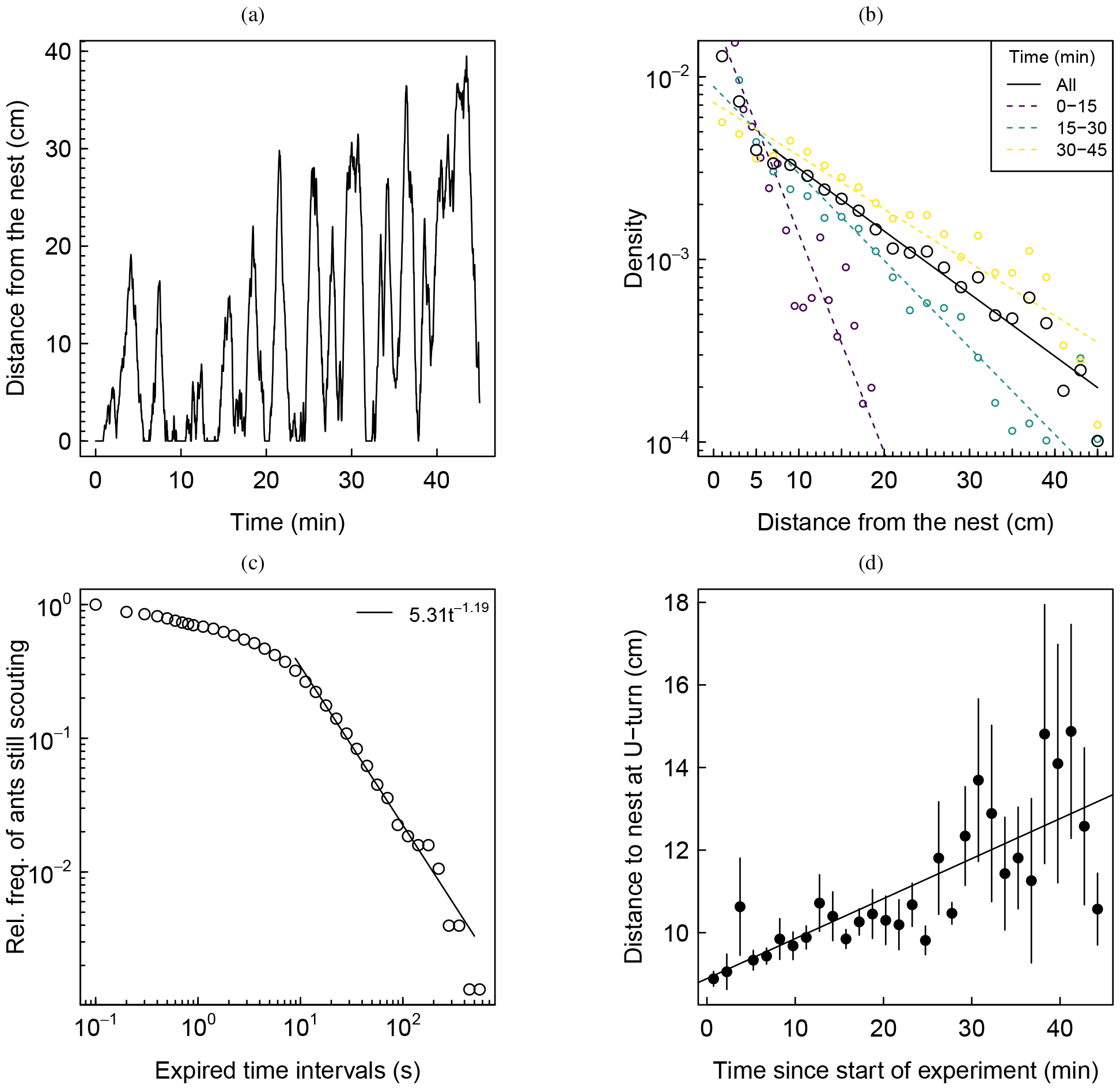
Analysis of the empirical data of [41] with *Temnothorax albipennis*. (a) Time series of the distance from the nest of an example ant for 45 min. The ant goes back and forth to its nest during the observation period. (b) Distribution of the distance from the nest of all ants (semi-log) across the entire observation period (black dots). Purple, green and yellow dots respectively show distributions for the first 15 minutes, from 15 to 30 minutes and from 30 to 45 minutes. Straight lines highlight the presence of exponential distributions. (c) Log-log plot of the survival curve of ants still in their outbound journey (scouting) before returning to their nest. For journeys longer than 1 s, logarithmic binning of the data is used, with bins 0.1 decade wide. Straight line shows the power law fit for outbound journeys longer than 10 s. (d) Maximum distance reached in searching journeys (mean*±*se) as a function of the elapsed time since experiment started. Mean and standard error calculated across 90-second bins. Straight line shows a linear model fit.

### 3.2 Simulations

We first simulated the original Boltzmann Walker model, a correlated random walk model without a mechanism for scouting agents to return to their nest. In this case, on average, agents progressively move away from their nest, as the number of reorientation events increases. The distribution of distances of agents – close to a Gaussian distribution, sees its mean and variance increase as the number of reorientation events increases (Figure 3a). This results in agents being more and more evenly spread in space – and further away from their nest, as the number of reorientation events increases. A typical measure of random walks resulting in diffusion is the Mean Square Displacement (MSD). As predicted for Brownian particles, the MSD of agents following the original Boltzmann Walker model increases linearly with the number of reorientation events, showing that agents are, on average, spreading steadily in all directions around their nest of origin (Figure 3b).

**Figure 3.**
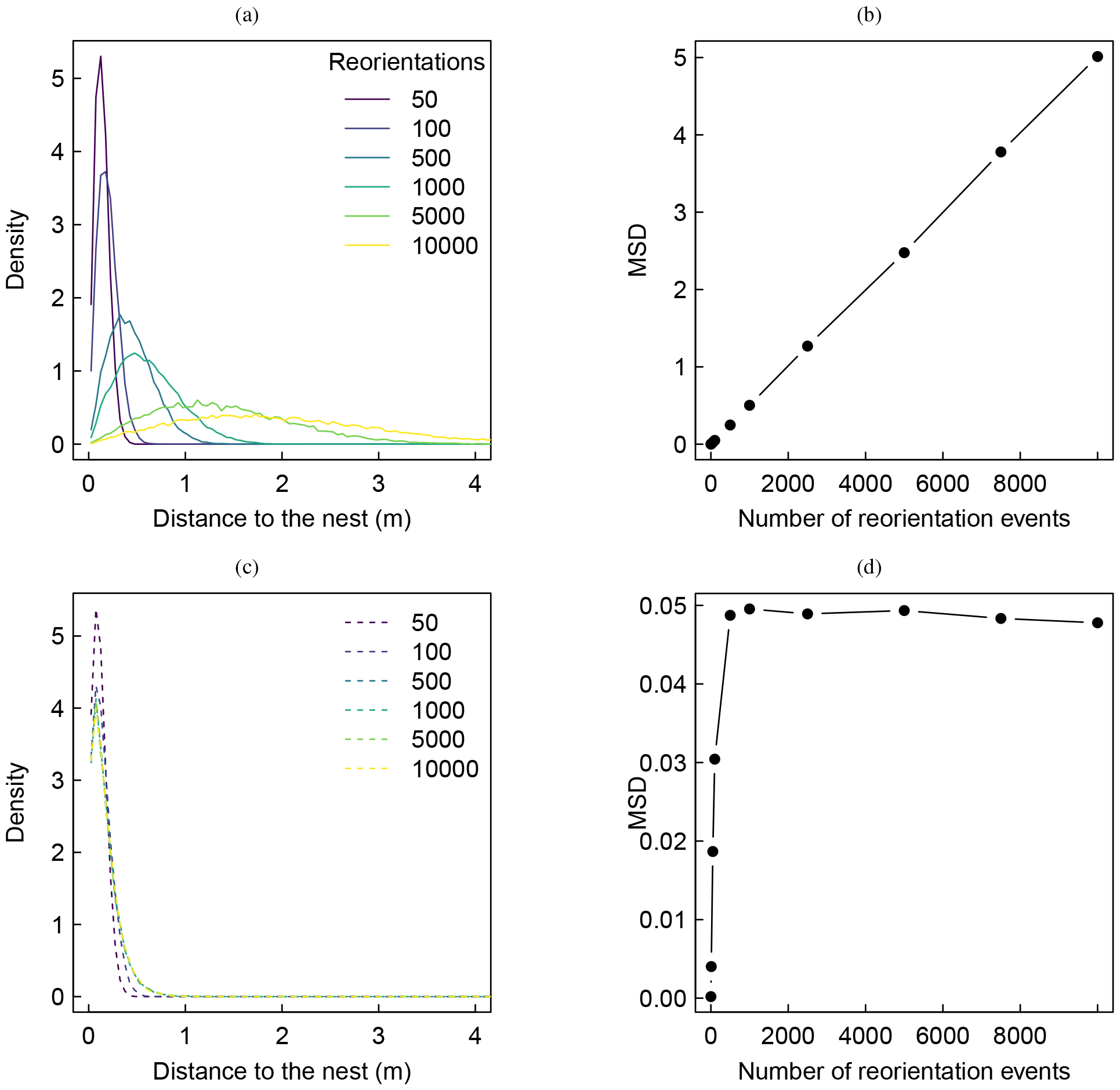
Distributions of distance of agents from their nests (a and c) and Mean Square Displacement (MSD, b and d) after a given number of reorientation events, without (a-b) and with (c-d) stochastic resetting. Random walks with stochastic resetting, with a probability to return to the nest constant with the number of reorientation events lead to a distribution of ants around their nest that is stationary, in contrast with the diffusive patterns observed in a random walk without resetting.

Introducing a mechanism for agents to return to the nest, by the means of a constant probability to return at each reorientation event changes this diffusing behaviour. After a few hundred reorientation events, the agents stop spreading steadily in space (Figures 3c-d). The distribution of distances to the nest does not change after 1000 reorientation events, quickly reaching a maximum spread (Figure 3c). On average, agents will therefore return to their nest before reaching the boundaries of this effective foraging area shaped by the introduction of a returning mechanism. This results in a MSD reaching a maximum after 1000 reorientation events (Figure 3d).

Our model of reference, in which the probability to return to the nest is constant across reorientation events (thereafter ‘model of reference’), predicts that for a given number of reorientation events, the distance of scouting agents from their nest is exponentially distributed (Figure 4a), in agreement with the empirical results found in laboratory experiments (Figure 2b). This is a key difference with the original Boltzmann Walker model: agents are now much more likely to be searching at short distances from their nests than in the Boltzmann Walker model, especially after a short period of time (a few reorientation events). After a longer period of time, the foraging range, at the colony-level, does not increase, with most of the scouting agents in the vicinity of the nest.

**Figure 4.**
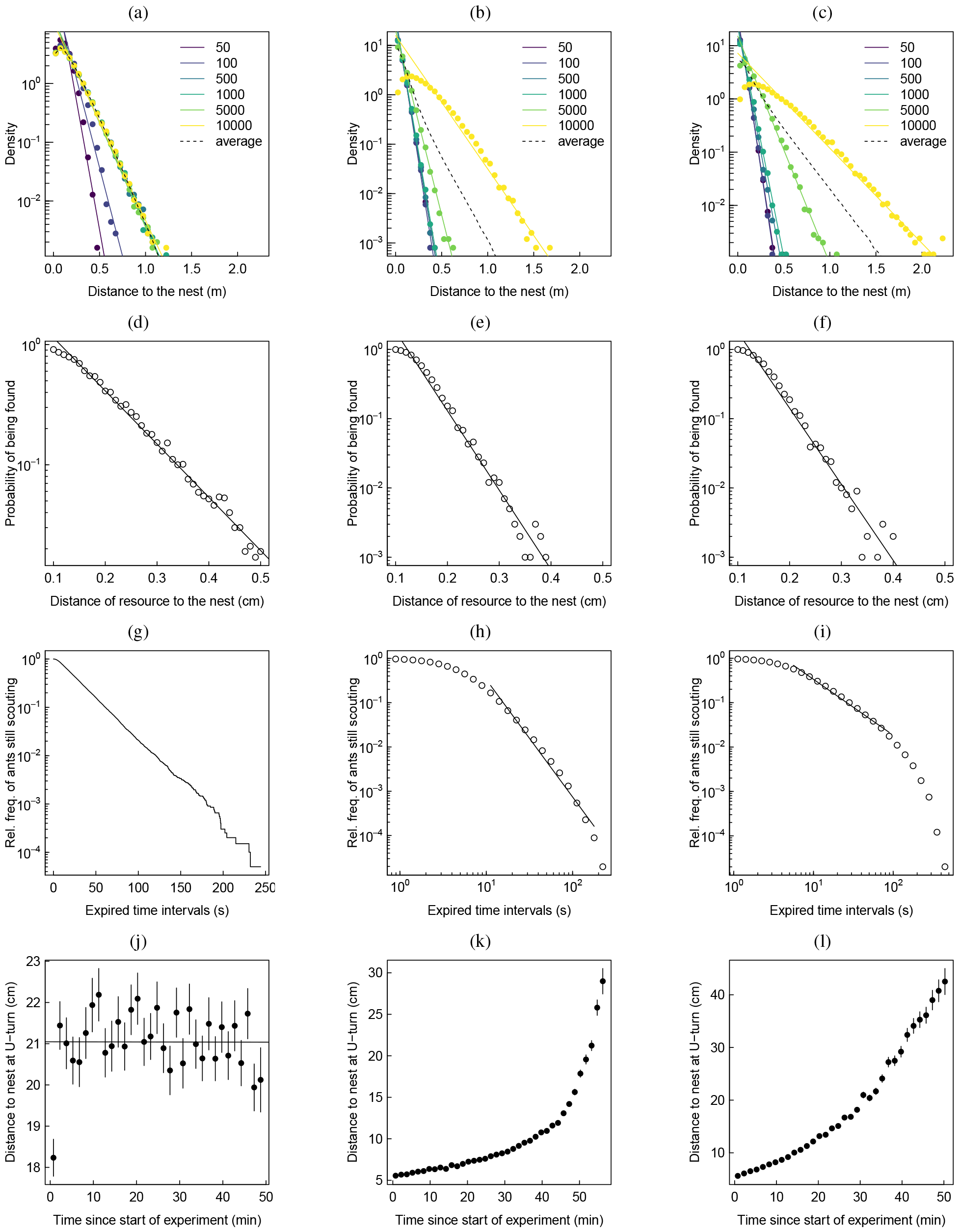
Distributions of distance of agents from the nest (a-c), Probability to find food as a function of distance (dashed line is averaged across all possible reorientation number conditions) (d-f), Survival curve (elapsed time without having returned to the nest), (g-i), and Maximum distance reached in a scouting bout as a function of time (j-l) for the three returning probability scenarios, constant with the number of reorientation events (column 1), or decreasing linearly (2) and exponentially (3).

We now turn to the comparison of three distinct implementations of the returning mechanism: that of reference, which is constant with the number of reorientation events, and two others, in which the probability to return to the nest decreases with the number of reorientation events, either linearly or exponentially (Figure 1). We find that all three implementations predict the distribution of distances of agents to their nest to be exponentially distributed (Figure 4a-c). It is however worth noting that once averaged over all number of reorientation events (i.e. from 1 to 10000) the distance distribution turns out to be clearly made out of two distinct exponential distributions (Figure 4b-c), similar to what we found in empirical data (Figure 2b). The general shape of the distribution (i.e. exponential) is the same across all returning mechanisms and is translated into the probability to find resources in the environment to also be exponentially distributed with the distance between the resource to the nest (Figure 4d-f). We investigated alternative intensities to return to the nest as a function of the number of reorientation events: decreasing step function and squared exponential and an increasing linear function (Figure S1). We find that all these alternative behavioural rules, including one in which the probability to return to the nest increases with the number of reorientation events, lead to scouts being exponentially distributed (Figure S2).

We also tested the effect of the underlying movement model in determining the exponential distribution of worker around their nest. We compared a pure random walk model (*g* = 0), in which agents do not have any direction of preference, and a ballistic random walk (*g* = 1), in which agents go straight forward in each of the reorientation events of their outbound bouts, to the correlated random walk model (*g* = 0.6) of reference (Figure S3). We also investigated the model of reference with path lengths uniformly distributed as opposed to exponentially distributed (Figure S4). All these different models lead to workers being exponentially distributed around their nest, leading to resource discovery probability exponentially decaying with distance to the nest. This suggests the density of workers being exponentially distributed around their nest to be an emergent property of a searching model in which individuals periodically return to their nest, irrespective of the movement patterns followed by the individuals.

There are however important differences between the three rules of returning to the nest. First, when the probability to return to the nest is not constant with the number of reorientation events, the emerging distribution stops being stationary (Figure 4a-c), at least until the probability to return reaches its minimum value. Second, there are major differences between the three alternative returning mechanisms regarding the survival curve of the scouting bouts (Figure 4g-i) and how the distance at which agents return to their nest changes with the duration of the searching behaviour (Figure 4k-l). With the constant probability of the model of reference, the survival curve is exponentially distributed (Figure 4g), in contrast to empirical results that suggest power-law distributions (Figure 2c and [27]). When the probability to return linearly decreases with the number of reorientation events, the survival curve looks power-law distributed for journeys longer that 10 s (Figure 4h). A similar pattern is observed with the probability to return decaying exponentially with the number of reorientation events, although the power-law relationship may not hold for journeys longer than 100 s (Figure 4i). Another important metric to discriminate the underlying behavioural rule is the relationship between the maximum distance from the nest reached during a bout and the time since the individual has started scouting. This metric provides further evidence that in *Temnothorax albipennis*, the probability to return to the nest is not constant with the number of reorientation events. While in empirical data we found that the maximum distance reached during outbound journeys is increasing with time (Figure 2d), agents with a probability to return to the nest constant with the number of reorientation events also have an average maximum distance reached during outbound journeys constant in time (Figure 4j). In contrast to this pattern, we find that rules in which agents have a probability to return to the nest decreasing with the number of reorientation events lead to agents reaching further distances away from their nest when the time since the start of their exploration increases (Figure 4k-l). We also find that both decreasing rules (linearly or exponentially decreasing) lead to slightly different relationships, with the linear decreasing giving the steepest non-linear increase of maximum distance reached as a function of elapsed time (Figure 4k-l).

## 4 Discussion

In this article, we propose a new model depicting the movements of searching and scouting ants, made of a discrete correlated random walk model and of a stochastic returning mechanism to the nest in the event of unsuccessful search. The main result of this study is that such a returning mechanism very robustly ensures that scouts exploring the surroundings of a nest will be exponentially distributed with distance from the nest. In other words, scouts will explore more efficiently the immediate surroundings of the nest than regions further away, no matter the actual rules governing their movements or their tendency to return to their nest.

In bees, it was recently proposed that individuals would have such a returning mechanism, but with a probability to return to the nest constant in time [20]. We show here, for a different movement pattern made of series of straight lines, that whether this probability is constant with the searching time or not actually does not change the shape of the emerging distribution of distance of scouts from their nest of origin. However, we stress here a major difference: when the probability to return to the nest is not constant in time, the emerging distribution is not stationary anymore, up to a point where the probability reaches its minimum value.

Our study further shows that, in ants, this probability to return to the nest is not constant with the number of reorientation events. We first show, analysing empirical data from a laboratory experiment of single *Temnothorax albipennis* ants searching a foraging arena, that the distance at which individuals initiate inbound journeys (homing) increases with time, leading to loops of increasing radius away from the nest, which is not compatible in a model with a probability to return constant in time. An additional evidence of these increasing loops is shown by the survival curve of searching bouts, which is not exponentially distributed but instead decreases as a power law in the empirical data with *Temnothorax albipennis* ants that we analysed, in contrast to the model in which the probability to return is constant. The fact that the survival curve of searching bouts was decreasing as a power law had already been reported in data collected in the field on fire ants [27]. The finding that searching ants make increasing loops around their initial location resonates with a behaviour reported in the field for desert ants of the genus *Cataglyphis* and *Melophorus bagoti* when searching for their nest entrance [23, 25]. In the Australian desert ant, it was also reported that the average distribution of the distance of searching ants from their initial location was made of two distinct exponential distributions in unfamiliar environments [25, 26], very similar to what we show in *Temnothorax albipennis*. The presence of two distinct exponential distributions was also reported in laboratory conditions in the ant *Aphaenogaster senilis* [28]. To explain these two exponential distributions, authors suggested the presence of two searching modes, either by the means of two distinct random walks [25] or with risk-prone and risk-averse strategies [28]. Here, our model and empirical data analysis suggest a new explanation: there is only one searching mode but the probability to return to the nest not being constant in time leads the distribution of distance to be made of two distinct exponential distributions once averaged across all ants. It is worth noting here that in our model the emergence of two exponential distributions appears only when averaging across all number of reorientation events. In other words, the two distributions are only visible when agents are not synchronised in their probability to return to the nest – i.e. when the considered population is made of agents whose probabilities to return to the nest are not the same. To conclude, we have multiple sources of evidence in searching and scouting ants of various species with different resource exploitation strategies (fire ants, ants from the genus *Cataglyphis*, Australian desert ants *Melophorus bagoti, Temnothorax albipennis* and *Aphaenogaster senilis*), in both field and laboratory conditions, of a returning mechanism made of increasing loops. Our simple model, made of a correlated random walk and stochastic returns to the nest (spatial resets) whose frequency decreases with time, predicts all the features reported in the literature previously and in the empirical data analysed for the purpose of our study – namely that searching ants are distributed around their nest following a distribution made of two exponential distributions and a survival curve of searching bouts distributed as a power law. This really highlights the relevance of random walk with stochastic resetting models to describe ant searching movements at both the individual and collective or colony levels.

Regarding the functions of the mechanisms that we found, this exponential distribution of distances may be a key mechanism at the colony-level to ensure that the close vicinity of the nests are more crowded that areas further away, improving defence and the ability of individuals to quickly alert the other nest members of any danger located at short distances from the nest [43]. At the collective level, the emerging spatial pattern of scouts around their nest, exponentially decreasing with distance to the nest, may therefore be the result of a trade-off between exploration of the environment and safety of the nest – and of the individuals. It may be that the simple mechanism of having a probability to return to the nest that decreases with time allows for the emergence of a composite search model, in line with theoretical predictions that showed that foraging bouts made of an intensive search phase, followed by an extensive phase if no food is found in the intensive phase, are efficient [44, 45]. However, the fact that we found that any movement mechanism combined with a returning mechanism will lead to an exponential distribution of distances questions whether this is an adaptive emerging pattern or rather a mandatory by-product of central place foraging in highly social species. Even if the general shape of the exponential distribution might not be adaptive, its actual parametrisation in different species might be. Different movement behaviours or returning rules may indeed well be finely-tuned so that the stationary distribution of scouts around the nest is well-suited to the needs and objectives of the colony.

Our model provides predictions at the nest-level while mainly requiring calibration from individual trajectories in simple and non-expensive laboratory experimental apparatuses. Our study suggests metrics at both collective and individual levels that may be suitable to investigate further the actual behavioural rules of movements and of returning used by different ant species, in relation to the pattern of scout distribution and exploration at the colony level. This avenue of research, in addition to shedding light on scouting behaviours and spatial patterns in social insects, may be of interest to improve estimates of the foraging area of a colony in the field, from simple laboratory experiments. Such estimates are indeed very costly and difficult to get in the field, requiring practitioners to track individuals and identify their colony of origin in complex environments. Our results regarding the density of scouts and defining a stationary foraging area are important for instance to understand better how colonies exploit resources in their environment – including the specific case of polydomous ants. Polydomous ants are ant colonies distributed across multiple spatially separated nests connected to each other and to foraging patches by a network of trails [46]. Predicting how scouts are distributed around their nest and, therefore, inferring the probability to find resources whether they lay within or outside the foraging area established by scouts with a returning mechanism is indeed instrumental to understand the costs and benefits of polydomy [47, 48] and the emergence of trail networks and polydomous networks, which are the results of colonies solving a quality-distance trade-off [4].

Our study, proposing a class of models with both spatial and temporal resets paves the way to investigate novel questions in the future. First, regarding the translation of the movement behaviours measured in the laboratory to estimates at the colony level in the field, it is unclear how accurately parameters estimated in laboratory experiments would describe behaviours occurring in nature. In other words, do ants adapt their returning behaviour when evolving under laboratory conditions in which their nest size and foraging area are heavily constrained or would this behaviour be finely tuned to natural conditions? Investigating this would be an important step in inferring foraging area from laboratory experiments. A second open question concern the time scale at which the returning probability operates. Since we showed, based on empirical observations, that this probability is likely not constant with time, and decreases with the duration of the searching journeys of ants, the question of the actual clock underlying this effect is still open. Moreover, it is unknown how asynchronous is this temporal reset in the scout population. Is this clock reset at the scale of the hour, when scouting workers start a new searching session? Or at the scale of the day, between resting periods? And how does the age of the worker interfere with this probability, with older scouts maybe having a lower probability to return on average, that would allow them to explore further away than younger ones – the colony age structure therefore affecting the foraging area? Different movement [22, 49] or returning rules coexisting may also underlie a complex stationary distribution made of several distinct exponential distributions. These questions are important to understand better the emergence of foraging or searching area in an unknown environment in ants.

## Supporting information

Electronic Supplementary Material

## Data availability

Hunt, Edmund R. et al. (2015). Data from: Ants determine their next move at rest: motor planning and causality in complex systems [Dataset]. Dryad. https://doi.org/10.5061/dryad.jk53j

## Footnotes

1 https://doi.org/10.5061/dryad.jk53j

