## Supplementary material for "Random walks with spatial and temporal resets may underlie searching movements in ants": Electronic Supplementary Material

Valentin Lecheval
School of Mathematics, University of Leeds, UK
Institute for Theoretical Biology, Department of Biology, Humboldt Universität zu Berlin, Berlin, Germany
Science of Intelligence, Research Cluster of Excellence, Berlin, Germany


Elva J.H. Robinson
Department of Biology
University of York, UK

Richard P. Mann
School of Mathematics
University of Leeds, UK

February 20, 2024

***Keywords*** resetting random walks · collective behaviour · animal movements · central place foraging · social insects

### List of Figures

S1 Three alternate probability intensities to return to the nest as a function of the number of reorientation events experienced by the agent. . . . 2

S2 Distribution of distances of scouts to their nest of origin (a-c) and probability to find resources as a function to their distance to the nest (d-f) for three alternate probability intensities to return to the nest, step function (a, d), increasing linear function (b, e) and squared exponential (c, f), as shown in Figure S1. 2

S3 Effect of the concentration of new directions, $g$, for three values of heading persistence: $g = 0$, leading to a pure random walk (a,d, g), $g = 0.6$, parameter of reference used in the main text standing for a correlated random walk (b, e, h), and $g = 1$ for null deviations, or complete persistence (c, f, i). a-c). Trajectories of a single ant for 1000 reorientation events, with parameters of reference and $g = 0$ (a), $g = 0.6$) (b) and $g = 1$ (c). Distribution of distances from the nest for scouts after 1000 reorientation events for $g = 0$ (d), $g = 0.6$) (e) and $g = 1$ (f). Probability to find a resource within 500 reorientation events as a function of its distance to the nest for $g = 0$ (g), $g = 0.6$) (h) and $g = 1$ (i) – calculated across 1000 searching ants for each distance. . . . 3

S4 Distribution of distances from the nest for scouts after 1000 reorientation events when lengths of paths are uniformly distributed, in contrast to being exponentially distributed (main text). Parameters of the uniform distribution are chosen so that the average path length is the same as in main text. Distribution of distances are shown for three distinct probability intensities to return to the nest, constant (a), linearly increasing (b) and exponentially decaying (c) with respect to the number of reorientation events. Straight line stands for a linear model fit and shows agreement to an exponential distribution. . 4

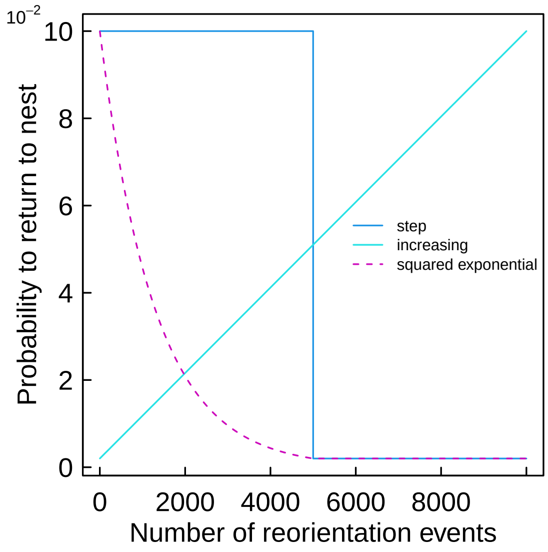


Figure S1: Three alternate probability intensities to return to the nest as a function of the number of reorientation events experienced by the agent.

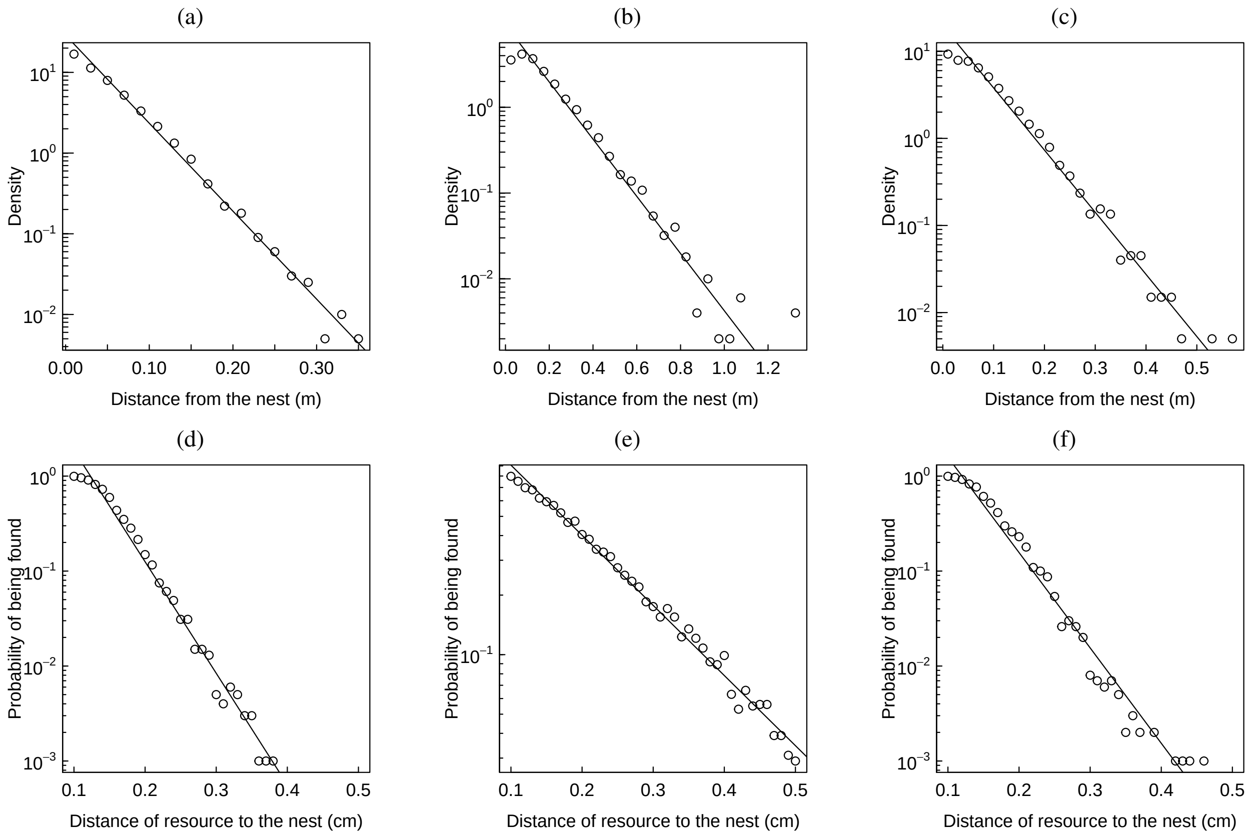


Figure S2: Distribution of distances of scouts to their nest of origin (a-c) and probability to find resources as a function to their distance to the nest (d-f) for three alternate probability intensities to return to the nest, step function (a, d), increasing linear function (b, e) and squared exponential (c, f), as shown in Figure S1.

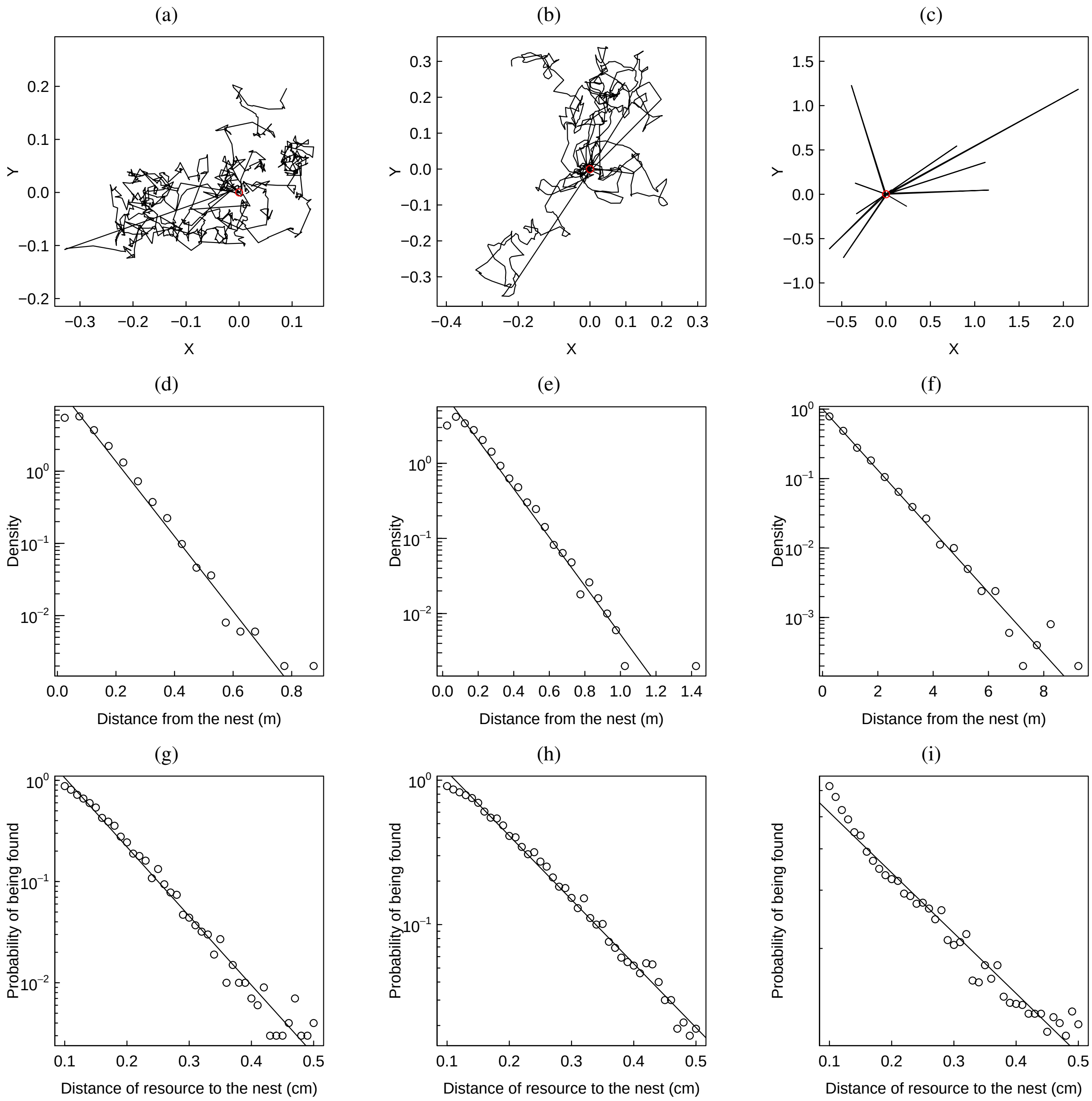


Figure S3: Effect of the concentration of new directions, $g$, for three values of heading persistence: $g = 0$, leading to a pure random walk (a,d, g), $g = 0.6$, parameter of reference used in the main text standing for a correlated random walk (b, e, h), and $g = 1$ for null deviations, or complete persistence (c, f, i). a-c). Trajectories of a single ant for 1000 reorientation events, with parameters of reference and $g = 0$ (a), $g = 0.6$) (b) and $g = 1$ (c). Distribution of distances from the nest for scouts after 1000 reorientation events for $g = 0$ (d), $g = 0.6$) (e) and $g = 1$ (f). Probability to find a resource within 500 reorientation events as a function of its distance to the nest for $g = 0$ (g), $g = 0.6$) (h) and $g = 1$ (i) – calculated across 1000 searching ants for each distance.

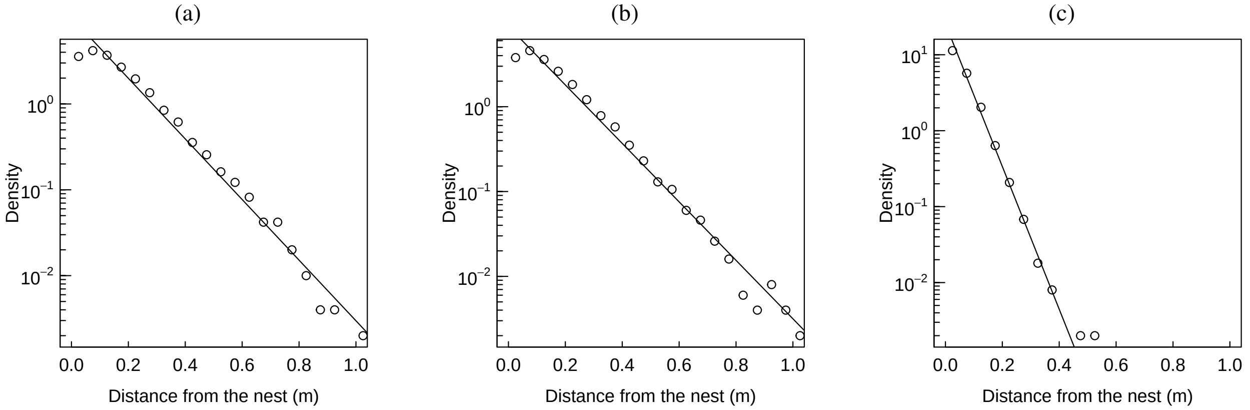


Figure S4: Distribution of distances from the nest for scouts after 1000 reorientation events when lengths of paths are uniformly distributed, in contrast to being exponentially distributed (main text). Parameters of the uniform distribution are chosen so that the average path length is the same as in main text. Distribution of distances are shown for three distinct probability intensities to return to the nest, constant (a), linearly increasing (b) and exponentially decaying (c) with respect to the number of reorientation events. Straight line stands for a linear model fit and shows agreement to an exponential distribution.
